# DNA Polymerase Beta Catalytic and Fidelity Mutations Drive Platinum Specific Drug Sensitivity

**DOI:** 10.64898/2026.06.15.732403

**Authors:** Jacob Lindquist, Josh Heyza, Istri Ndoja, Nasrin Movahhedin, Chris Yunker, Marina Cardo-Vila, Seongho Kim, Joann Sweasy, Steve M. Patrick

## Abstract

With the advent of genome sequencing and its widespread use in the clinic, there is a great need to identify mutational biomarkers that predict therapeutic responses. DNA polymerase Beta (Polβ) and the base excision repair (BER) pathway have been previously implicated as modulators of response to platinum-based chemotherapies and are mutated in as high as 30% of cancers. Here, we show in a triple-negative breast cancer (TNBC) model that two classes of mutations in Polβ, reduced catalytic activity (E295K and D256A mutation) and reduced fidelity (I260M), are sufficient to drive cisplatin and carboplatin-specific sensitivity. Cellular response to oxaliplatin in these Polβ mutant models is minimal relative to cisplatin and carboplatin. Additionally, we show that sensitivity is associated with reduced repair of both platinum-induced DNA intrastrand adducts and interstrand crosslinks (ICLs). Downregulation of the upstream BER factor uracil DNA glycosylase (UNG) reverses drug sensitivity consistent with these Polβ mutations negatively impacting ICL DNA repair to drive drug sensitivity. In addition, intrastrand adduct repair readout indicates these lesions also play a role in the sensitivity observed in Polβ mutant models. *In vivo* studies demonstrate a significant effect on tumor growth delay with cisplatin treatment in tumor xenografts harboring Polβ mutations. These results support the potential for using Polβ mutations as predictive biomarkers for cisplatin and carboplatin therapies in the clinical setting.

## Introduction

Platinum-based chemotherapies are among the most commonly used treatments in cancer, treating patients with cancer types such as ovarian, lung, head and neck, and breast tumors (1–3). These drugs target and bind to DNA, resulting in inhibition of DNA replication and transcription which can lead to apoptosis (4). However, recurrence and resistance to these drugs are commonly seen in the clinical setting, highlighting the need to better predict patient response. Multiple mechanisms of cisplatin resistance have been previously described, including decreased drug accumulation, increased drug inactivation, as well as alterations in DNA repair (5). Increased DNA repair has been shown in cell culture models as well as in patient response to cisplatin, and it is seen in nearly all cases of high cisplatin resistance (4).

Platinums directly bind to DNA resulting in the formation of mono-adducts, intrastrand adducts, as well as interstrand cross-links (ICLs) (6,7). These adducts distort the DNA, leading to fork stalling during DNA replication and transcription and eventual apoptosis if the cell is unable to process or repair these platinum-induced adducts. The main DNA repair pathway that resolves intrastrand adducts is the nucleotide excision repair (NER) pathway, which can also work in conjunction with the homologous recombination (HR) and Fanconi Anemia (FA) pathways to remove ICLs (8,9). Unsurprisingly, increased activity of these DNA repair pathways has been associated with chemoresistance to cisplatin (10). Despite understanding the major pathways that repair platinum-DNA adducts, the impact of other pathways as well as various clinically relevant mutations in additional DNA repair factors on cisplatin adduct repair is incompletely understood.

Although all platinum reagents are able to bind to DNA, the intrastrand adducts and ICLs formed by oxaliplatin versus cis/carboplatin differ in the complex structures and distortions induced in the DNA double helix. In particular, the ICLs formed by cis/carboplatin at GpC sites and the covalent link between guanine bases on complementary DNA strands result in the flanking cytosines being extrahelical and flipped away from the duplex structure which is distinct from the ICLs formed by oxaliplatin (11–16). Loss of BER as well as mismatch repair (MMR) drives a cis/carboplatin resistance, and these two pathways work in an epistatic manner to mediate cis/carboplatin response (17). The extrahelical cytosines adjacent to the ICL produced by cis/carboplatin can be de-aminated by specific APOBEC3 family members, after which uracil DNA glycosylase (UNG) is able to generate an abasic site (17). AP endonuclease (APE1) is then able to cleave the 5’ end of the abasic site, allowing for recruitment of polymerase β (Polβ) to incorporate nucleotides. In the case of nucleotide misincorporation by Polϐ, downstream MMR MutSα (MSH2-MSH6) can bind and continue to block ICL DNA repair. Overall, this process results in non-productive processing of the platinum ICL, resulting in persistence of ICL lesions and sensitivity to cis/carboplatin (17–19).

Polβ is the main DNA polymerase involved in short-patch BER and is mutated in up to 30% of tumors including lung, colon, gastric, and breast cancers (20, 21). Several of these mutations have been previously characterized and shown to trigger cellular transformation as well as induction of both resistance and sensitivity to platinums based on varying phenotypes (22–24). Previous knockdown of Polβ was shown to induce a cisplatin-resistant phenotype both in *in vitro* and *in vivo* models, while expression of a catalytic-deficient mutant form of Polβ in these same models induced a platinum hypersensitivity (17–19, 24). Previous work suggested that APOBEC3 family member expression is an important variable that modifies how loss of specific Polβ mutations impact platinum-based chemotherapy responses, especially in regard to the impact on ICL non-productive repair (17–19).

Breast cancer remains the most commonly diagnosed and second most lethal cancer amongst woman in the U.S. (25). In part this is due to the triple-negative subtype (TNBC), which unlike other subtypes, still lacks targeted therapies. Recent research has highlighted a promising role for platinum use in the treatment of TNBC (26). However, there remains a large heterogeneous response (27), highlighting the need for predictive biomarkers to help determine the best treatment option. This need for predictive biomarkers alongside the heterogenous response to platinum amongst various Polβ mutations highlights the need for further investigation into specific Polβ mutations that drive platinum sensitivity. In this study, we investigate the ability of catalytic-deficient Polβ to induce hypersensitivity to cis/carboplatin. We show through multiple mutations (E295K, D256A and I260M), that catalytic deficiency as well as reduced Polϐ fidelity results in platinum sensitivity in a cis/carboplatin specific fashion. Importantly, this work highlights the contribution of both intrastrand adducts and ICLs in driving the sensitive phenotype in these Polβ mutant models. These results highlight the potential role of specific Polβ mutations in modulating platinum DNA adduct repair highlighting their potential utility as predictive biomarkers for predicting platinum response in the clinical setting.

## Materials and Methods

### Cell Culture

All cells were maintained at 37°C in a 5% CO_2_ incubator. MDAMB231 cells were grown in RPMI (Dharmacon) containing 10% fetal bovine serum (FBS) and 1% penicillin/streptomycin (Dharmacon). SUM1315 cells were grown in F12/DMEM containing 10% FBS and 1% penicillin/streptomycin. Cells were tested for Mycoplasm contamination and used within 10 passages for all experiments.

### shRNA transfection and Stable Knockdown Generation

Cells knocked down for WT Polβ and re-expressing mutant D256A were generated previously (provided by the Robert Sobol lab)(18,19). Other knockdowns were generated via Mission shRNA plasmid DNA. Lentiviral particles were generated using HEK293T cells using the following plasmids: PMD2G, PMDLG/RRE and PRSV/RRE. Lipofectamine 3000 (Invitrogen) was used to transfect plasmid DNA. 24hr post-transfection, the media was changed to media of target cells. Viral particles were harvested and centrifuged 72hr post-transfection before filtration through 0.2-μm filters. Viral stocks were aliquoted and used fresh or stored at −80°C. Polybrene was added to viral media immediately prior to transfection to maximize efficiency. Transfected cells were selected and stable knockdowns were maintained using puromycin (Sigma-Aldrich) or G418 (Sigma-Aldritch).

### Quantitative Real-Time PCR

Cells were harvested and pelleted before extracting RNA via TRIzol (Invitrogen) as previously described (17). RNA concentration was quantitated using a nanodrop spectrophotometer and cDNA was derived from 2ug of RNA using the High-Capacity cDNA Reverse Transcription Kit (Applied Biosystems). Real-time PCR was performed in technical triplicate using PowerUP SYBR Green Master Mix (Applied Biosystems), with GAPDH as an endogenous control. Relative gene expression was determined with the 2ΔCT method and performed in biological triplicate. Primer sequences were as follows:

**Table 1.**
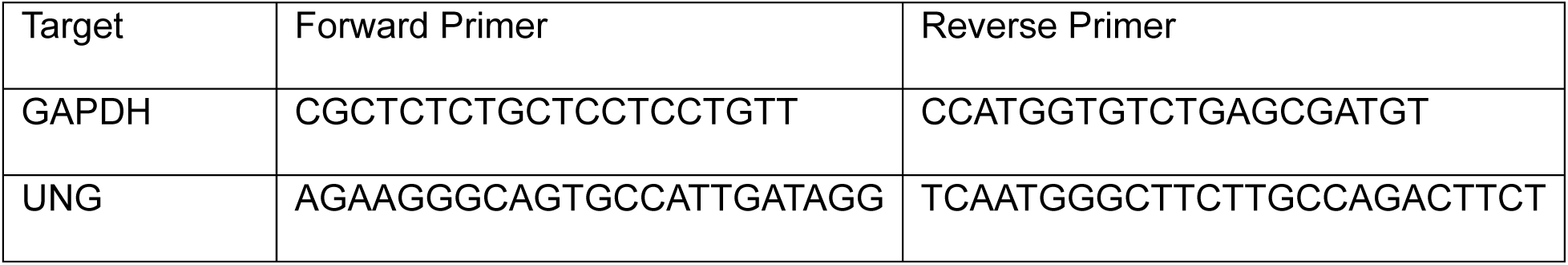
RT-PCR Primers.

### Colony Survival Assay

300-800 cells were seeded into 60-mm dishes one day prior to treatment. Cisplatin (Sigma-Aldritch), carboplatin (Sigma-Aldritch), and oxaliplatin (Sigma-Aldritch) stock concentrations were prepared at 1mM in phosphate-buffered saline (PBS). Once drug was completely dissolved via vortexing, solutions were filtered through a sterile 0.2-μm filter. Cisplatin and carboplatin stock solutions were prepared fresh for each experiment. Oxaliplatin and methyl methanesulfonate (MMS, Sigma-Aldrich) stock solutions were stored at −80°C and -20°C, respectively, following preparation and used within a 6-month period. For cisplatin, carboplatin, and oxaliplatin, cells were incubated with varying concentrations for 2hr in serum-free medium. For MMS, cells were treated in serum-containing medium for 4 hours. Following treatment, drug media was removed and replaced with complete medium and colonies were allowed to grow for 7 days. Plates were stained and fixed with crystal violet solution (20% ethanol, 1% w/v crystal violet). Plates were counted for colonies with ≥50 cells and percent survival was determined using the ratio of the number of colonies in treated plates compared to the average number of colonies in 0 concentration plates multiplied by 100. Experiments were performed in biological and technical triplicate. IC50 values for each replicate were calculated with effect values at each concentration using CompuSyn.

### MTS Viability Assay

**∼**2000 cells were seeded per well into a 96-well plate one day prior to treatment. Cells were treated with cisplatin for 2 hours in serum-free medium at working concentrations diluted from stock solutions prepared as for colony assays. After treatment, drug medium was removed and cell viability was measured 48 hours post-treatment. For MTS assays, 10uL of respective reagent was added to each well and allowed to incubate at 37°C for 2-4 hours. Plates were read at 450nM (MTS) and percent survival was determined as a ratio compared to the average reading at 0 concentrations.

### Modified Alkaline Comet Assay for Cisplatin Interstrand Crosslinks

Modified alkaline comet assay was used to analyze the presence and repair of ICLs as previously described (17–19). Cells were treated with cisplatin (10μM) for 2 hours and collected at 0, 24, 48, and 72 hours post treatment. Immediately prior to collection, cells were incubated with fresh hydrogen peroxide (100 μM) for 15 minutes to induce DNA strand breaks or control buffer. The use of hydrogen peroxide to induce strand breaks serves as a way to fragment DNA and is a comparative control in measuring ICLs present. The presence of ICLs covalently links the DNA strands, thus retarding migration through the gel (17–19). Once cells were collected, suspensions containing ∼10,000 cells were embedded in 0.5% low melting point agarose and pipetted onto a slide pre-coated with 1% agarose. Slides were incubated at 4°C in lysis buffer (2.5 M NaCl, 10mM Tris, 100 mM EDTA, pH10, 1% Triton X-100) for 1hr. After lysis, slides were incubated at 4°C in alkaline electrophoresis buffer (300 mM NaOH, 1 mM EDTA, pH > 11) for 20 min. Electrophoresis was carried out at 300 mA and 20–25 V for 25 minutes. Slides were neutralized in a neutralization buffer (0.4 M Tris–HCl, pH 7.5) for 10 mins before DNA precipitation in 95% ethanol for 10 min. DNA was stained with SYBR Gold (Invitrogen) and comets were imaged using a Nikon fluorescence microscope. At least 50 cells were imaged and analyzed per slide using ImageJ and OpenComet v1.3 plugin. Data are expressed as comparing the mean olive tail moment of cisplatin plus hydrogen peroxide with hydrogen peroxide-treated and untreated control samples, which represents the percentage of ICLs remaining on the fragmented DNA (17,18).

### Slot-blot Assay for Intrastrand Adduct Repair

Following cisplatin treatment and time course for DNA repair, cells were collected and genomic DNA was isolated. DNA (2μg) from corresponding cells was loaded into each well and vacuumed onto a nitrocellulose membrane. The membrane was incubated at 80°C for 30 minutes, followed by blocking solution (5% milk in PBS-T) for 1hr and incubated overnight with α-Pt-GG antibody (1:1000, MABE416). The following day, the membrane was washed in PBS-T buffer four times for 15 min each before incubation with α-Rat secondary for 1hr. Membranes were washed for 5min with 2mL of combined ECL reagent before being imaged on ChemiDoc imaging system (Biorad). Densitometry values of each blot were determined using ImageJ. Briefly, values for each band were measured and background-adjusted before standardizing to relative 0 time points.

### In Vivo Study

36 female athymic NUDE mice (six mice per group) were purchased for this study from Envigo and were maintained in accordance with protocols approved by the Wayne State University Institutional Laboratory Animal Care and Use Committee. The mice were allowed to acclimate for 1 week before handling. Tumors were established by implanting 5 x10^6^ cells subcutaneously into the rear flanks of each mouse. Weight of the mice was measured regularly starting on the day of implantation and through the experimental endpoint. Once tumors became visible, volume was measured with calipers and defined as (width^2^ × length/2). After tumors reach an average volume of ∼100mm, mice were randomly sorted into groups and treated with either vehicle (PBS) or cisplatin (3 mg/kg) by intraperitoneal injection at the indicated times. Mice were sacrificed once the burden of the tumor reached ∼1000 mm^3^.

### Vector Generation for CRISPR Knock-In of I260M

DNA oligonucleotides were purchased from IDT and used to clone the sgRNA into BbsI-digested pX330-U6-Chimeric_BB-CBh-hSpCas9 (a gift from Feng Zhang (Addgene plasmid # 42230) (28). The homology-directed repair template was cloned by Gibson Assembly into pFastBac Dual. Homology arms and a codon-optimized sequence containing the Polβ I260M mutation and the remaining C-terminal coding sequence was purchased from IDT as G blocks. These G blocks, a PCR amplified puromycin resistance cassette, and linearized pFastBac Dual backbone were cloned together by Gibson assembly. Gibson reaction product was then transformed into chemically competent bacteria and plasmid isolated using a Qiagen MiniPrep kit. Correct assembly was validated by Sanger sequencing. A similar approach was utilized for Polβ E295K generation.

### CRISPR-Cas9 generation of Knock-in Clones

I260M knock-in cells were created via vectors generated in-house. Briefly, cells were transduced with pFastBac Dual vector containing the homology-directed repair sequence to knock-in the I260M mutation and a puromycin resistance cassette (downstream of the Polϐ stop codon) as well as px330 containing Cas9 and the sgRNA. Three days post-transfection, cells were selected with puromycin before isolating clones. Once clones grew to adequate size, genomic DNA was harvested from cells and used alongside PCR primers flanking the genomic location of incorporation.

E295K knock-in cells were generated by the Sweasy lab similar to the approach used for I260M knock-in generation and DNA sequence verified to be homozygous knock-in in MDAMB231 cells. Briefly, cells were lysed in Lamelli buffer + ß-mercaptoethanol, scraped and boiled for 15 min. at 90°C. Protein fractions were analyzed by 10% SDS-PAGE and transferred to PDVF membrane. The membrane was then blocked with a 50-ml solution of 5% nonfat milk, 1× Tris-buffered saline (TBS; 50 mM Tris–HCl, pH 7.4, 150 mM NaCl), and 0.1% Tween 20 (TBS-T) for one hour at RT. After washing the blot twice with 1× TBS-T, primary antibody of Polβ (at a dilution of 1:1,000; Abcam cat# ab26343) was added in a 5-ml solution of 5% nonfat milk and 1× TBS-T, and the blot was incubated overnight at 4°C. Subsequent to being washed with 1× TBS-T solution three times, the membranes were washed twice with 1× TBS. The membranes were then incubated with anti-rabbit horseradish peroxidase-conjugated secondary antibody (at a dilution of 1:25,000, cat#NA9340V GE) for 1 hr at RT. The membranes were then washed as described above. Proteins were visualized using an enhanced chemiluminescence kit used according to the manufacturer’s directions (ThermoFisher). Actin-HRP antibody (Sigma cat#A3854) at 1:10,000 in 5% BSA-TBS-t for 30 min, was used to normalize the protein loading. The levels of proteins were quantified by normalizing the levels to the housekeeping protein using Image Lab software (BioRad).

### Statistical analysis

All experiments were performed with a minimum of three technical and biological replicates unless noted otherwise. Data was summarized with means and standard deviations. Where appropriate, distribution of data was checked for normality and a log transformation was applied where needed. Unpaired t-tests were used to distinguish differences between two groups and one way ANOVA followed by post-hoc analysis was used for groups of three or more. All data analyses were carried out using the EZR package (R commander Version 2.8-0, R version 4.2.2, RStudio Inc., Boston, MA) and SigmaPlot version 10.0.

## Results

### Sensitivity of Polβ catalytic Mutants to Cis/Carboplatin

Our previous studies have demonstrated that knockdown of Polβ and other BER/MMR members led to a platinum resistant phenotype while the knockdown of endogenous Polβ and re-expression of a catalytic dead Polβ mutation (D256A) showed a cisplatin sensitive phentotype (17). We sought to assess and compare another catalytic dead Polβ mutation, E295K, that was previously shown to have catalytic deficiency in a triple-negative breast cancer model (24). A CRISPR-Cas9 knock-in approach was taken to homozygously express this mutation in MDAMB231 TNBC cells. These cells were confirmed for deficiency in BER activity by treating with alkylating agent Methyl Methanesulfonate (MMS) (**Supplemental Figure 1A**). To determine if this cisplatin-sensitive phenotype was a result of catalytic mutant Polβ, we first compared sensitivity to cisplatin (**Figure 1A**) and found similar sensitivity in both the D256A and E295K Polβ mutations. Additionally, we measured cell viability for these cells in an MTS assay treated with cisplatin and found a similar trend (**Supplemental Figure 1B**). In accordance with prior research, we expected similar sensitivity to carboplatin due to the adducts formed being functionally identical to those induced by cisplatin. In these Polβ mutant models, we observe similar degrees of carboplatin sensitivity as compared with cisplatin (**Figure 1C**). In contrast, the ICLs induced by oxaliplatin do not form extrahelical cytosines to the same extent as cis/carboplatin, and thus, would not be expected to be recognized by APOBEC3 members which induce cytosine deamination to uracil that are subsequently processed by the downstream BER pathway (17). Thus, we did not initially expect to observe oxaliplatin sensitivity in these Polβ mutant models assuming drug sensitivity is driven solely by the non-productive processing of ICLs. Interestingly, we observed a modest enhanced oxaliplatin sensitivity in both the D256A and E295K Polβ mutant models (**Figure 1E**). These results suggest there are additional effects/processes potentially including blocking intrastrand adduct repair that may be impacted by catalytically deficient Polβ mutations that drive the drug responses in these models.

**Figure 1.**
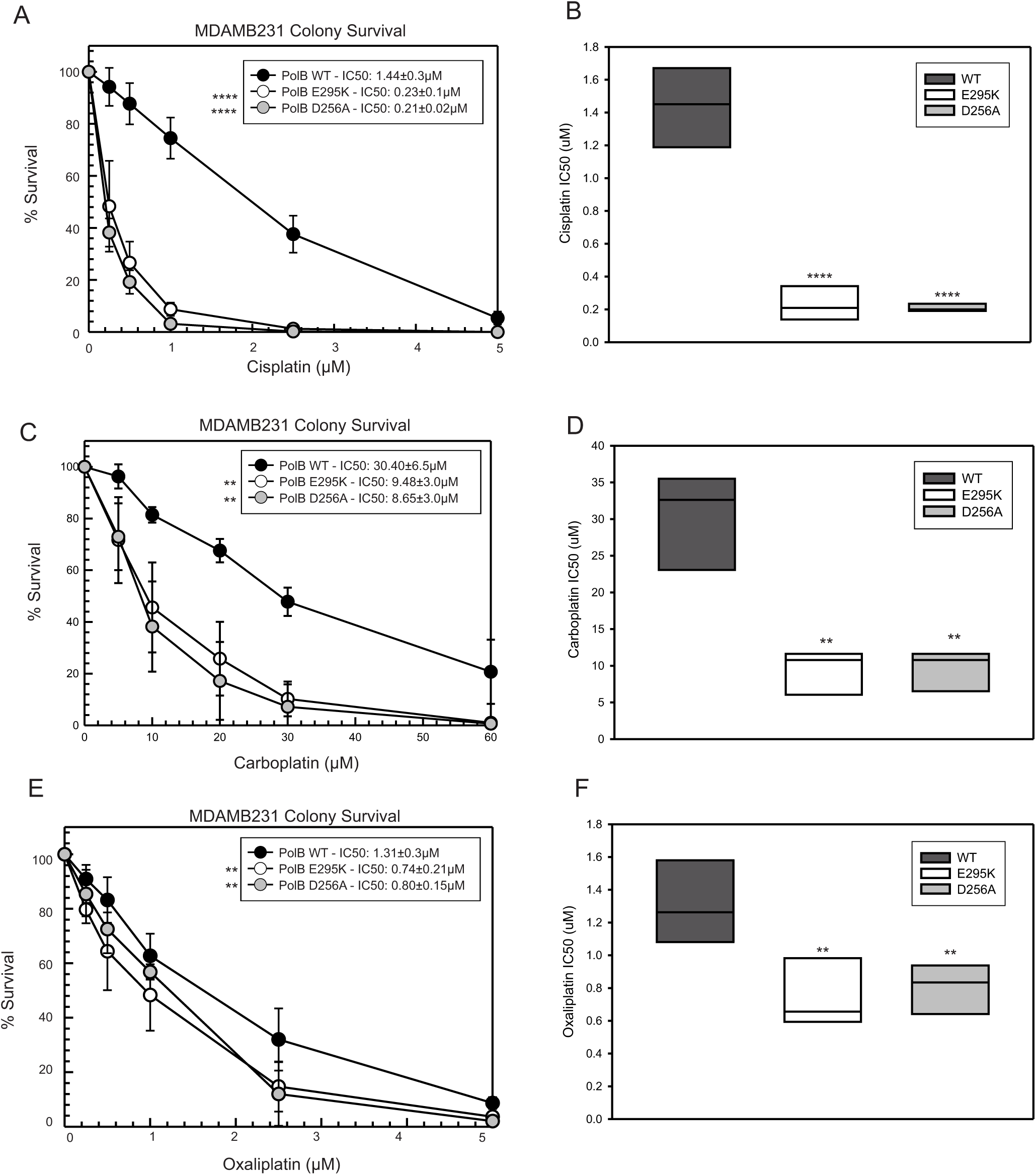
Catalytic Deficient Polymerase Beta induces selective platinum sensitivity. Colony survival assay of MDAMB231 cells treated for 2 hours with A) cisplatin, C) carboplatin and E) oxaliplatin. IC50 values plotted for cisplatin (B), carboplatin (D) and oxaliplatin (F). Data plotted as average ± SD of three independent replicates. IC50 values were calculated with CompuSyn software using a best fit of effect values at each dose and is presented as mean ± SD. IC50 values were compared to WT cells via a two-tailed t-test (* p<0.05, **p<0.01, ***p<0.001, ****p<0.0001).

### Reduced Rate of Adduct Repair in Polβ Catalytic Deficient Clones

In previous BER knockdown experiments, we found loss of this pathway to be associated with an increased rate of ICL DNA repair relative to wildtype (WT) expressing cells (17–19). Therefore, we analyzed the rates of repair of the cisplatin-induced ICLs in our catalytic deficient Polβ expressing cells via modified comet assays. We observed that the increased sensitivity is associated with a reduced rate of repair of the cis/carboplatin ICLs (**Figure 2A**). The accumulation and formation of ICLs peaks at ∼18-20 hours and thus, in all cases there is an increase in ICLs from the time 0 to 24hr timepoints followed by repair of the ICLs from 24 to 72 hr timepoints (17–19). At 24hr, 48hr and 72hr post treatment, there is a significant retention of ICLs in the Polβ E295K and D256A mutant cells compared with WT cells demonstrating inefficient ICL DNA repair (**Figure 2A**). Previously published results demonstrated that the loss of BER pathway proteins had no significant impact on the repair of cisplatin induced intrastrand adducts (18). In these Polβ mutant models, however, utilizing an intrastrand adduct specific slotblot assay (antibody targets the primary GG intrastrand adduct), we demonstrate that the catalytic deficient Polβ mutants also result in reduced rates of intrastrand adduct repair following cisplatin treatment (**Figure 2B,2C**). These data suggest that catalytic deficient Polβ drives cisplatin sensitivity through inhibition of both ICL and intrastrand adduct DNA repair.

**Figure 2.**
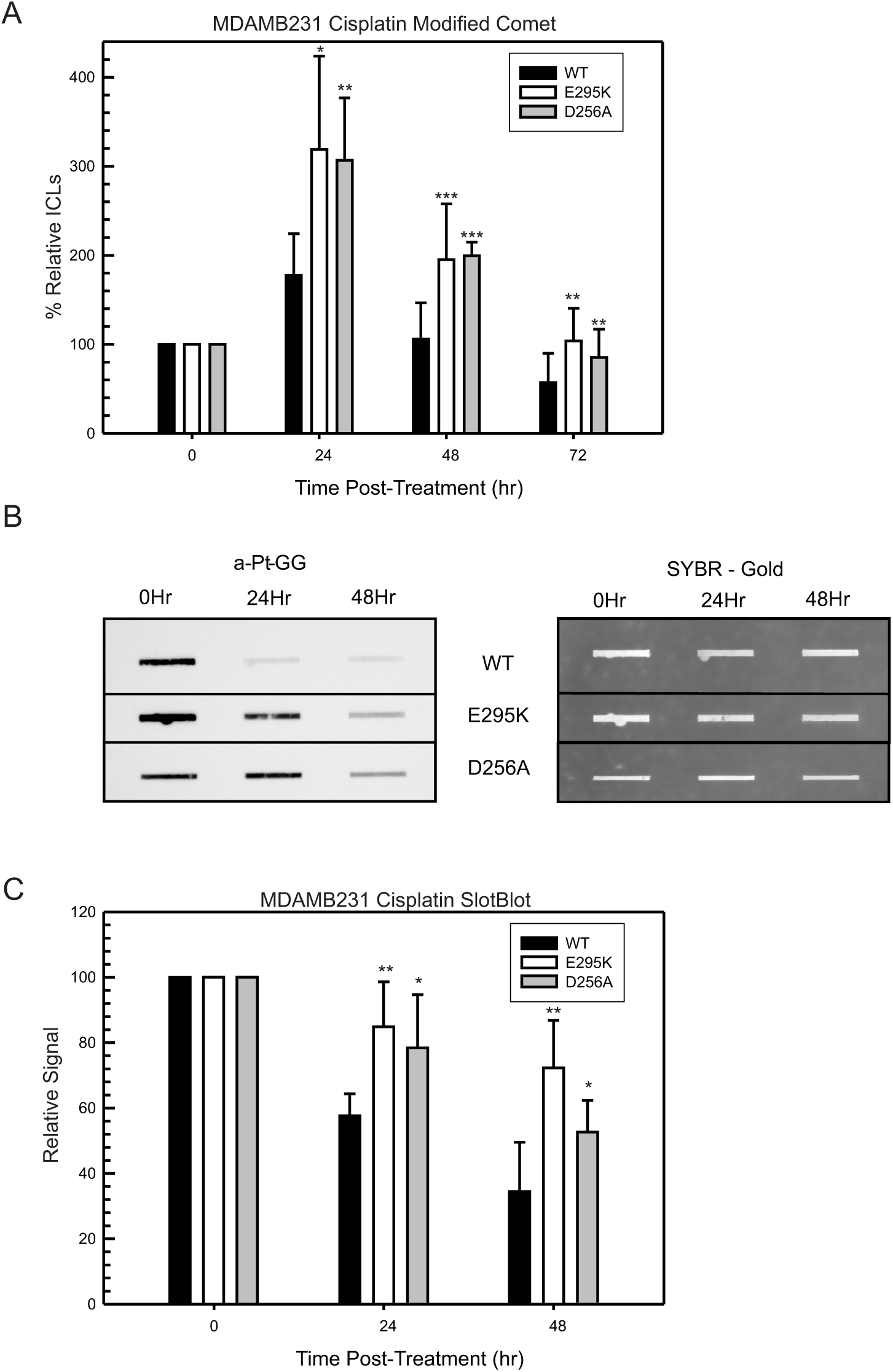
Both Intrastrand adduct and ICL repair are inhibited in Polβ mutant expressing cells. A) Modified Comet assay of cells treated with cisplatin (10μM) for 2 hours. Data is standardized to 0 hour post-treatment time points and presented as mean ± SD of three replicates. B) Slotblot assay against platinum GG intrastrand adducts of DNA collected from MDAMB231 cells treated with cisplatin (10μM) for 2 hours. C) Band quantification of slotblot assay using ImageJ software. Data is standardized to 0hour post-treatment time point and is present as mean ± SD of three independent replicates. P-values were determined relative to WT sample using a two-tailed t-test (* p<0.05, **p<0.01, ***p<0.001, ****p<0.0001).

### Inhibition of Upstream BER Partially Restores Cis/Carboplatin Resistance

We showed previously that loss of BER resulted in a cisplatin resistant phenotype that was correlated with an increased rate of ICL DNA repair (17–19). Here, to further define the mechanism(s) driving therapeutic response, we targeted multiple members of the BER pathway by shRNA knockdown and chemical inhibition. The results show resistance to cisplatin in WT MDAMB231 triple-negative breast cancer cells similar to what we previously observed (**Figure 3A**, WT vs shUNG)(18,19). If the drug sensitivity observed in the catalytic Polβ mutant models is solely attributed to the non-productive processing of ICLs, then knockdown of UNG would result in a complete reversal of the sensitive phenotype. The UNG knockdown experiments result in only partial reversal of the sensitivity of the Polβ catalytic E295K (**Figure 3A**, Ctrl vs KD) expressing cells as well as those with D256A mutation (**Figure 3B**, Ctrl vs KD) to cis/carboplatin. To ensure that incomplete reversal of cisplatin sensitivity was not due to insufficient UNG knockdown, we assessed transcript levels of UNG via RT-qPCR and found >90% knockdown in both WT and E295K expressing cells and >65% knockdown in D256A expressing cells (**Figure 3C**). This lack of complete resistance rescue was substantiated further with inhibition of APE1 incision by methoxyamine (MX) (**Supplemental Figure 2A**) as well as utilizing knockdown of APOBEC3 family members by shRNA (**Supplemental Figure 2B,2C)** for both Polβ catalytic mutant models. To determine if there were effects on the repair of ICLs, we used the modified comet assay and found that knockdown of upstream UNG resulted in increased rates of ICL DNA repair in both WT Polβ expressing cells (Ctrl vs UNG), as well as E295K expressing cells (Ctrl vs UNG) (**Figure 4A**). UNG is upstream of Polβ recruitment and thus, if non-productive ICL DNA repair contributes to the sensitivity of these catalytic Polβ mutant models, then downregulation of UNG would increase ICL DNA repair. These results are consistent with that concept and highlight that non-productive ICL processing is contributing to some of the enhanced sensitivity in these Polβ mutant models. As expected, downregulation of the BER factors had no impact on the repair of intrastrand adducts as determined via slotblot assay in both WT expressing cells (Ctrl vs UNG) and E295K expressing cells (Ctrl vs UNG) (**Figure 4B,4C**). These data are consistent with previously reported experiments where loss of BER pathway members was observed with only an increased rate of ICL repair (17–19).

**Figure 3.**
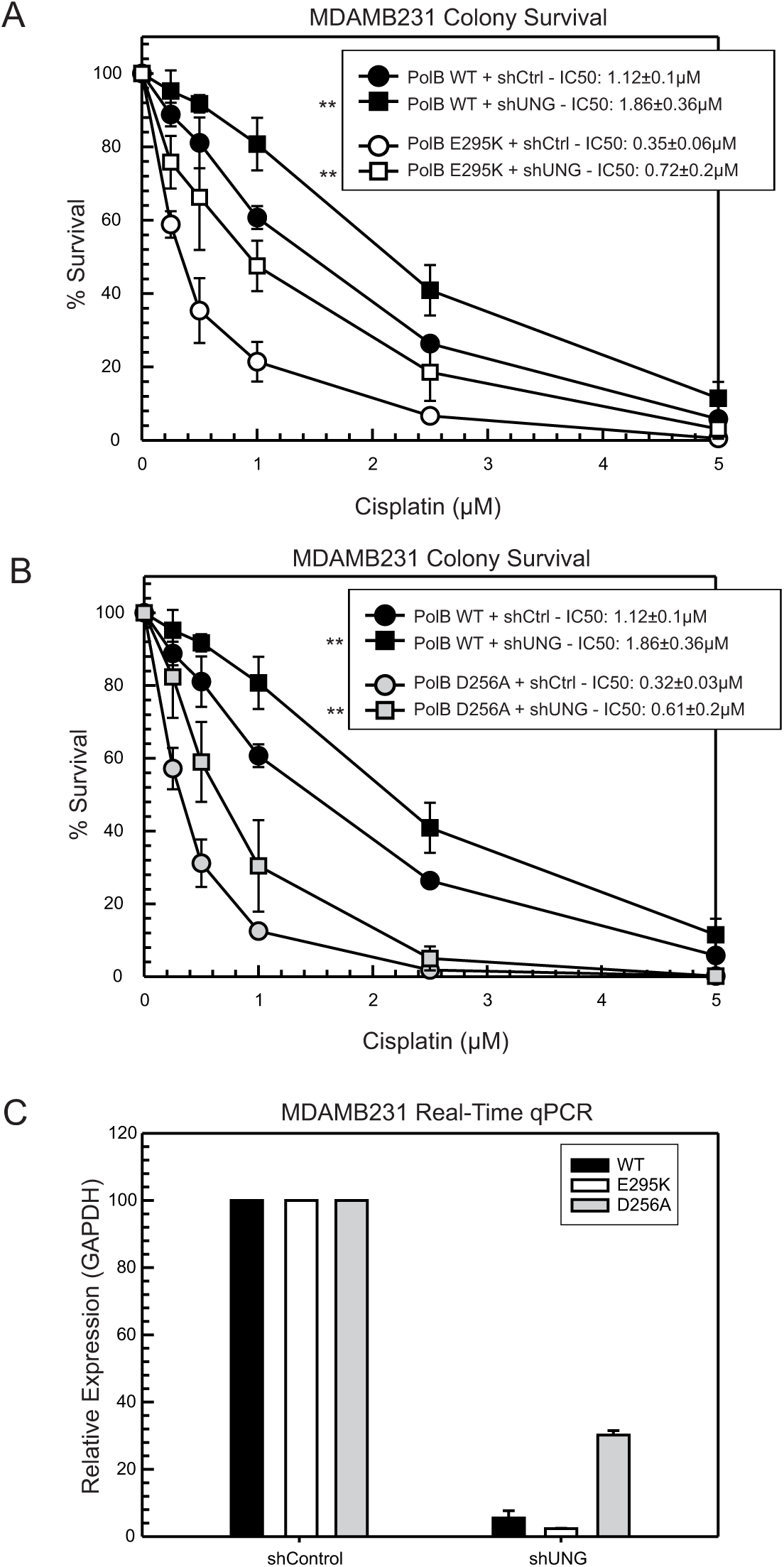
Inhibition of upstream UNG only partially induces resistance in catalytic deficient Polβ cells. Colony survival assay of cells expressing either shCtrl or shRNA targeting upstream UNG were introduced in Wild-Type or A) E295K and B) D256A expressing cells and treated for 2 hours with cisplatin in serum-free media. C) Realtime-qPCR of UNG in shControl and shUNG expressing cells. Relative expression was acquired with GAPDH relative control and expressions were standardized to shControl values. Data plotted as average ± SD of three independent replicates. IC50 values were calculated with CompuSyn software using a best fit of effect values at each dose and is presented as mean ± SD. IC50 values and relative expression of shUNG were compared to WT via a two-tailed t-test (* p<0.05, **p<0.01, ***p<0.001, ****p<0.0001).

**Figure 4.**
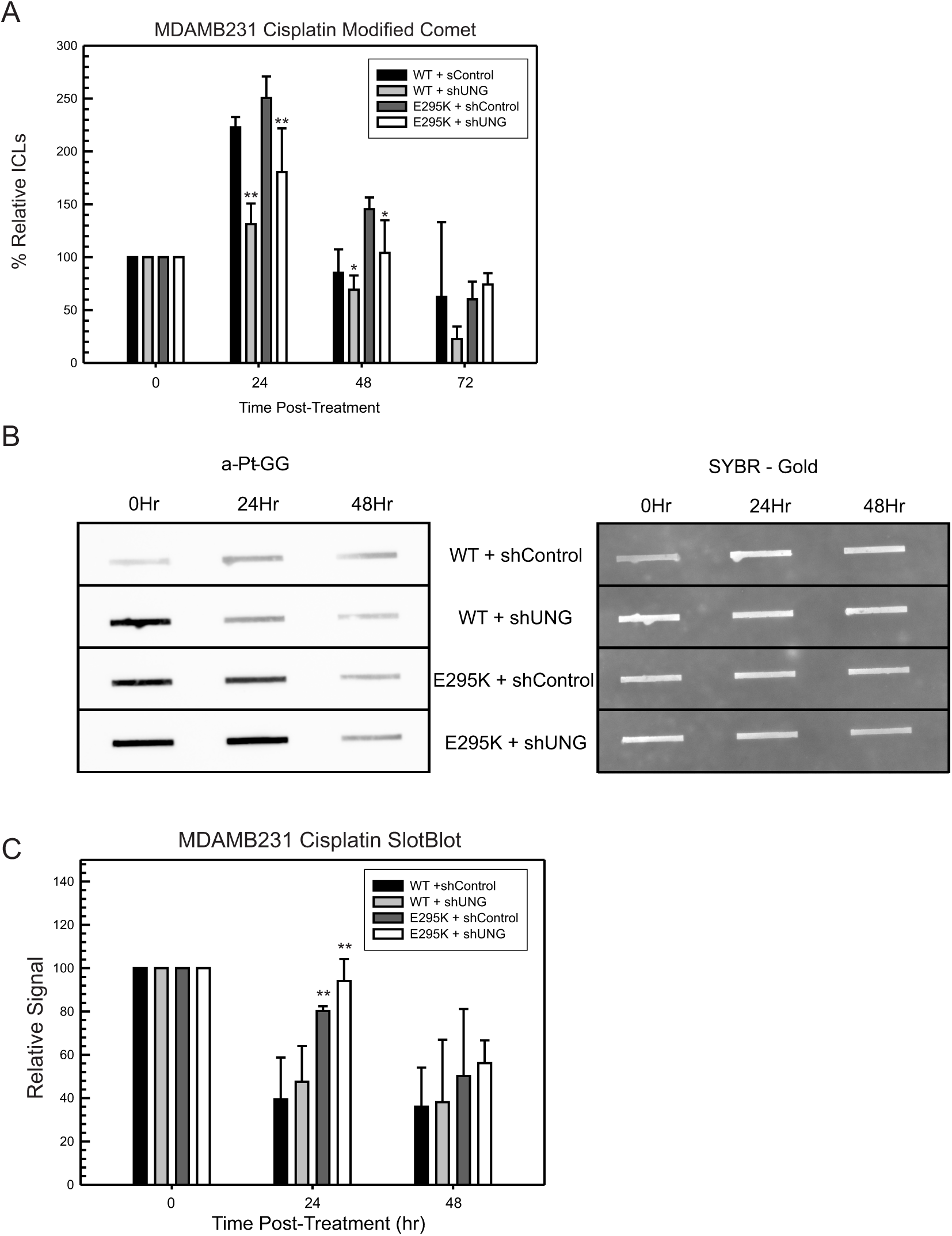
Inhibition of UNG results in increased ICL repair by not intrastrand adduct repair. A) Modified comet assay of WT and E295K cells expressing either shControl or shUNG were treated with cisplatin (10μM) for two hours and assessed at 0, 24, 48, and 72 hours post-treatment. Data was standardized to 0 hour time point and plotted as average ± SD of three independent replicates. B) Slotblot assay of WT and E295K expressing cells with shControl and shUNG were treated with cisplatin (10μM) for 2 hours and assessed for Pt-GG intrastrand adducts at 0, 25, and 48 hours post treatment. SYBR gold was used as a loading control. C) Band quantification of slotblot assay using ImageJ software. Data plotted as average ± SD of three independent replicates. Comparisons at each time point were compared via a two-tailed t-test (* p<0.05, **p<0.01, ***p<0.001, ****p<0.0001).

### Reduced Fidelity Polβ Clones Also Display Platinum sensitivity

To gain further clinical implications, we tested another Polβ mutant (I260M) in our cell-line models. The I260M mutation is associated with some polymerase catalytic deficiency as well as a mutator phenotype based on its reduced fidelity (*i.e.*, more likely to insert the incorrect nucleotide) (29,30). To accomplish this, we first looked at the expression of Polβ in several breast cancer cell lines as seen in **Supplemental Figure 3**. To incorporate this mutation, we utilized a CRISPR-Cas9 knock-in methodology. The advantage of this was the ability to insert a puromycin selection cassette to enrich for edited cells. We isolated clones from puromycin-resistant cell lines, isolated genomic DNA, and screened for knock-in by PCR (**Supplemental Figure 4**). Of note, we were unable to identify clones with homozygous knock-in of the I260M mutation. There are several potential underlying reasons for this, one being that the expression of this mutant in a homozygous manner is too toxic for cells to tolerate. Previous work has shown other mutations of Polβ present in a dominant negative fashion (24,29), highlighting the overall impact that a single allelic variant can have for this protein. Next, we tested several of these clones for cisplatin and carboplatin sensitivity and found similar levels of sensitivity as compared with the E295K and D256A expressing cells when expressed in the MDAMB231 triple negative breast cancer cell line model (**Figure 5A, 5B**). We also managed to knock-in the I260M variant in another TNBC cell line model (SUM1315) and found that this mutation also sensitized these cells to cisplatin and carboplatin compared to WT cells (**Figure 5C,5D**), however, the drug sensitivity was at a lower fold difference relative to MDAMB231 cells. One likely reason for this reduced fold difference in sensitivity is that SUM1315 express Polβ at lower levels relative to the MDAMB231, thus having a lower amount of Polβ to inhibit the repair of cis/carboplatin induced adducts (**Supplemental Figure 3**). These data support that both catalytic and fidelity Polβ mutations can inhibit the repair of cisplatin and carboplatin specific adducts and drive sensitivity to these drugs.

**Figure 5.**
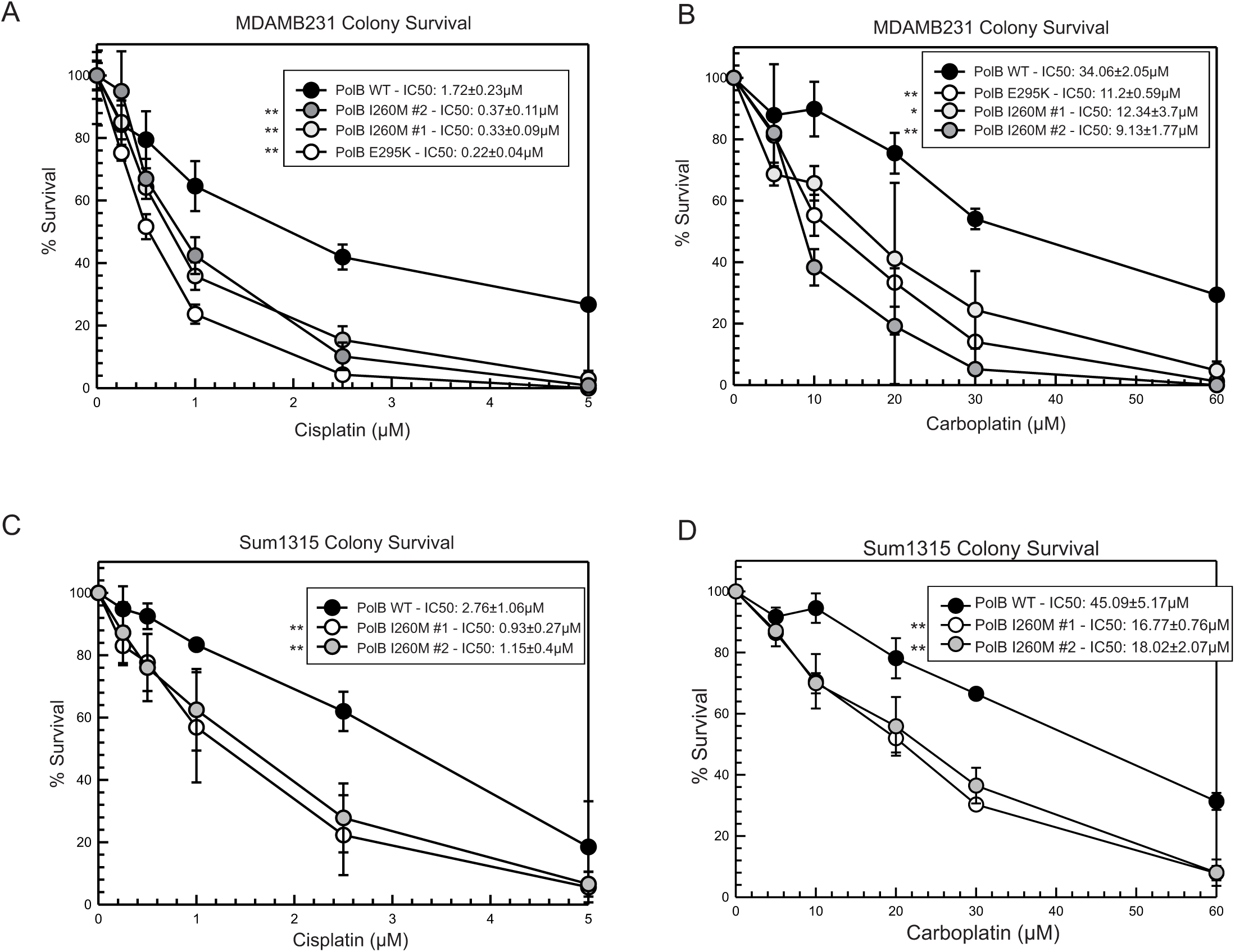
I260M induces cisplatin and carboplatin sensitivity in MDAMB231 and SUM1315 cells. Colony survival assay of MDAMB231 cells expressing either Polβ WT, E295K, or CRISPR knock-in clones expressing I260M variant were treated for 2 hours with A) cisplatin or B) carboplatin. Colony survival assay of SUM1315 cells expressing either Polβ WT or CRISPR knock-in clones expressing I260M variant were treated for 2 hours with C) cisplatin or D) carboplatin. IC50 values were calculated with CompuSyn software using a best fit of effect values at each dose and is presented as mean ± SD. IC50 values were compared to WT cells via a two-tailed t-test (* p<0.05, **p<0.01, ***p<0.001, ****p<0.0001).

### Polβ Catalytic Deficient Clones Induce Tumor Growth Delay With Cisplatin Treatment

Based on our previous *in vitro* data suggesting that catalytic deficiency of Polβ results in cisplatin sensitivity, we assessed whether these mutations would result in greater tumor growth inhibition in an *in vivo* mouse model. To accomplish this, we implanted MDAMB231 cells (WT, D256A, or E295K) subcutaneously into the flanks of nude mice to establish xenograft tumor models. Once tumors were established (∼75mm), we randomly assorted mice into either vehicle-or cisplatin-treated groups (controlling treatments and cell-type for average tumor size) and followed the treatment scheme outlined in **Figure 6A**. Briefly, mice were treated with either saline-solution or cisplatin solution (3mg/kg) via intraperitoneal injection. Mice were monitored for body weight loss during the experiment and displayed no signs of toxicity. The mice harboring tumors from catalytic deficient Polβ cells were found to have a significant tumor growth delay compared to those with WT expressing cells and harbored significantly smaller tumors at the termination of the study (**Figure 6B,6C**, WT vs E295K vs D256A). This data provides clear evidence that cisplatin-based therapies could be potentially beneficial to those patients harboring tumors with catalytic deficient Polβ.

**Figure 6.**
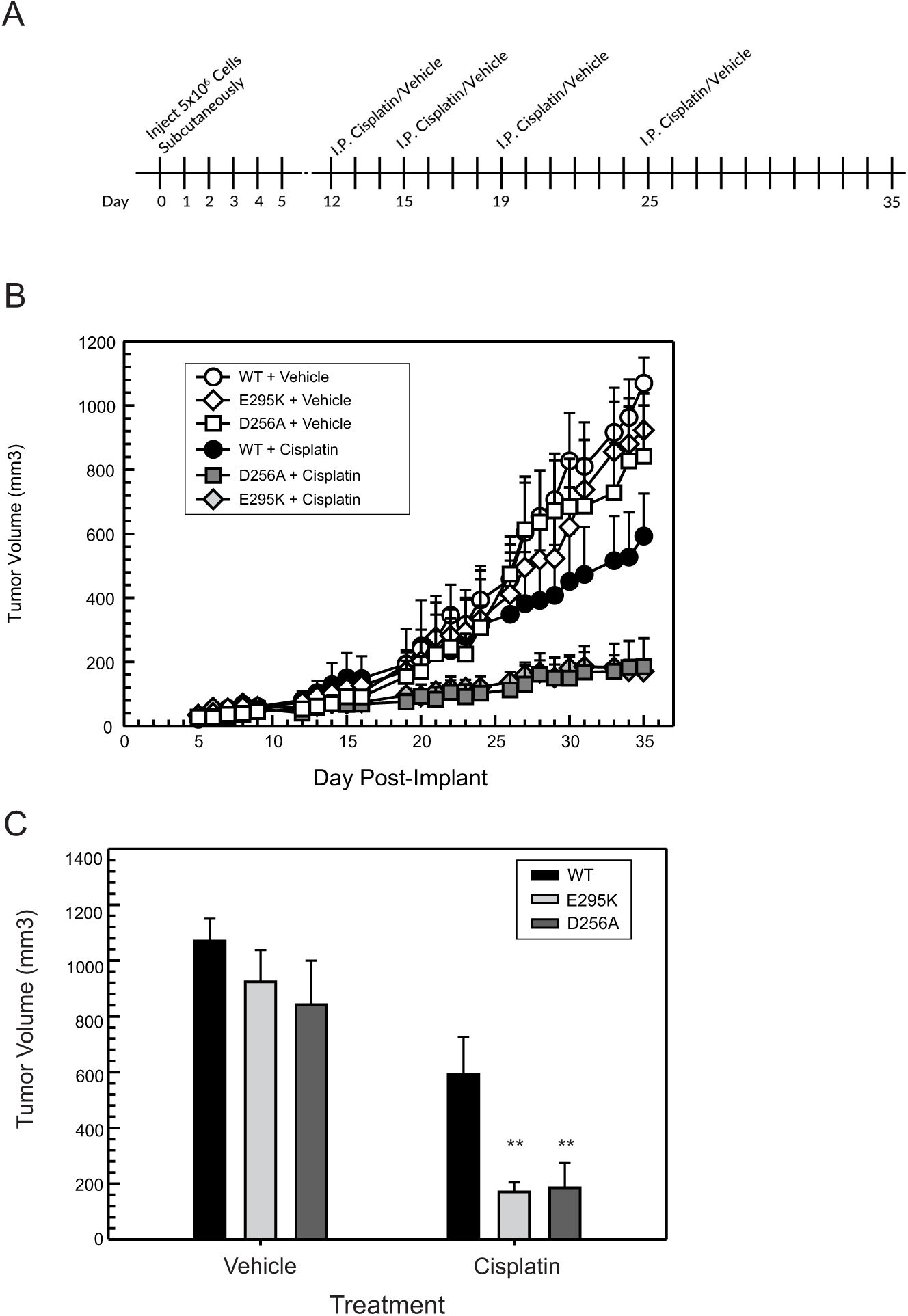
Catalytic Deficient Polβ drives a robust response to cisplatin in an *in vivo* mouse xenograft model. Mice were implanted with either WT, E295K, or D256A expressing cells and treated with vehicle or cisplatin. A, treatment scheme for the study. B, Tumor volume was monitored and final tumor volumes were reported over time. Data were plotted as the average ± SD of each group (6 mice per group). C, Final tumor volumes were compared using a two-way ANOVA and post-hoc test (* p<0.05, **p<0.01).

## Discussion

In this study, we show a cis/carboplatin specific sensitivity induced by several mutations of Polβ. These mutations affecting polymerase catalytic activity and fidelity led to a reduction in the repair of both intrastrand adducts as well as ICLs (**Figure 7; proposed mechanistic models**). We show that this increased sensitivity is partially impacted by downregulation or inhibition of upstream BER factors that would normally induce cis/carboplatin resistance via non-productive ICL processing. We specifically show that these Polβ mutations inhibit the repair and removal of both intrastrand and ICLs. Additional work has shown that an increased catalytic phenotype mutant D160G also results in an increased cisplatin sensitivity (31). This increased sensitivity was also associated with a reduced rate of repair in intrastrand adducts, in part through inhibition of XPA binding. Interestingly, the P242R Polβ mutation which is associated with partial loss of catalytic activity, results in cisplatin resistance (22). The authors show that the P242R Polβ mutation results in an XPA interaction which likely stimulates NER to repair the intrastrand DNA adducts (22).

**Figure 7.**
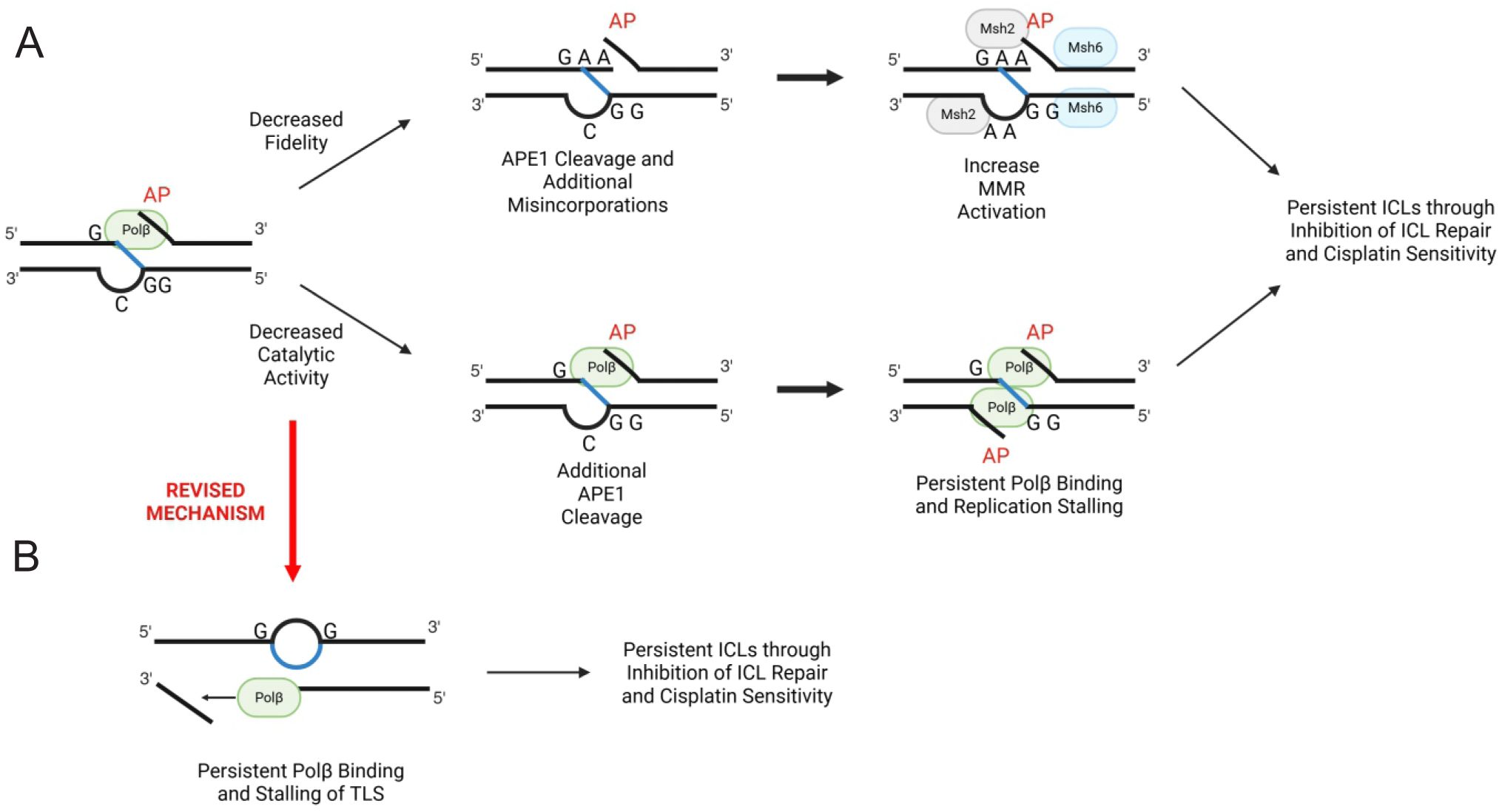
Proposed Mechanistic Model of Mutant Polβ Sensitivity to Cis/Carboplatin. The cells expressing mutant Polβ are shown to be selectively sensitive to cisplatin and carboplatin treatments. We show that this sensitivity is associated with a reduced rate of both intrastrand adducts and ICL repair induced by these drugs. A, In regard to the ICL response, we hypothesize that Polβ is able to bind to these ICL sites as a consequence of APOBEC3 family member deamination of extrahelical cytosines (17). The deamination of cytosine to yield uracil flanking the ICL is subsequently removed by UNG. APE1 is recruited and incises 3’ of the ICL to release the abasic site (AP) which enables Polβ recruitment. Catalytic deficient Polβ mutations can stall the polymerase at the ICL and block productive repair while Polβ fidelity mutants can result in introduction of the wrong nucleotides at the ICL site and drive more mismatch repair (MMR) protein which similar to catalytic mutants can lead to persistent ICLs and cis/carboplatin sensitivity. In B, both catalytic and fidelity Polβ mutations can also impact the translesion synthesis (TLS) past an intrastrand adduct and also block NER access and repair of these lesions to contribute to cis/carboplatin drug sensitivity.

These results highlight the importance of investigating the various clinically relevant Polβ mutations to fully understand which can mediate the response to platinum-based chemotherapy. To fully appreciate the potential clinical usage of Polβ mutations in being considered in genomic screening assays for predicting platinum-based chemotherapy response, more studies need to be conducted. There is some additional work reviewing the possibility of machine learning in helping to determine which mutations would be useful in predicting treatment courses (32). The variety of mutations and ultimate loss or gain of phenotype in the different domains of Polβ drive the need for continued research into the mechanism of how this protein works in multiple settings. The variety in phenotypes also suggest that crystallography of Polβ-DNA interactions with the addition of a platinum adducts could be useful in determining residues critical at these sites. Overall, our data suggests that Polβ interacts non-productively at both cisplatin intrastrand and ICL DNA adducts, helping to drive a sensitive phenotype.

The xenograft *in vivo* data highlight the potential tumor response to cisplatin in these Polβ mutant models. The significant tumor growth delay in these Polβ mutant models further highlight the potential use of these mutations in the clinical setting to help predict and drive better patient responses to specific types of therapy, including cis/carboplatin. Treatment for patients consists of several weeks and is primarily used in a neoadjuvant setting in stage 4 TNBC (33). Loss of Polβ has shown to be a driver in mutagenesis and tumorigenesis (24,30,34). Thus, targeting patients whose tumors harbor Polβ catalytic or fidelity mutations with multiple rounds of platinum therapy should lead to more robust and durable tumor responses and lead to better overall patient survival.

