## supplemental data for "DNA Polymerase Beta Catalytic and Fidelity Mutations Drive Platinum Specific Drug Sensitivity"

**Supplemental Figure 1: Polβ catalytic deficient mutants results in BER Deficiency and Display Sensitivity to Cisplatin.** (A) Colony survival assay of MDAMB231 cells expressing either WT Polβ or E295K variant were treated for 4 hours with alkylating agent MMS in serum free media. (B) MTS viability of MDAMB231 cells treated with cisplatin for 2hrs and assessed 72hours later. Data plotted as average ± SD of three independent replicates. IC50 values were compared to WT via a two-tailed t-test (* $p<0.05$, **$p<0.01$, ***$p<0.001$, ****$p<0.0001$).

**Supplemental Figure 2: Inhibition of upstream APE1 and APOBEC3 only partially induces resistance to cisplatin.** (A) Colony survival assay of MDAMB231 cells expressing either WT Polβ or E295K variant were pre-treated with 20μM of APE1 inhibitor methoxyamine for 4 hours before treatment with cisplatin for 2hours in serum-free media. (B and C) Colony survival assay of MDAMB231 cells expressing either shCtrl or shRNA targeting Apobec3 members were introduced in Polβ Wild-Type and B) E295K and C) D256A expressing cells. Data plotted as average ± SD of three independent replicates. IC50 values of shUNG were compared to shControl via a two-tailed t-test (* $p<0.05$, **$p<0.01$, ***$p<0.001$, ****$p<0.0001$).

**Supplemental Figure 3: Protein Expression of Polβ in breast cancer cell lines.** Protein was collected from breast cancer cell lines and western blot conducted using an anti-Polβ antibody and β-Actin loading control.

**Supplemental Figure 4: PCR Demonstrates Incorporation of I260M Allele via Homology Template.** Genomic DNA from individual cell colonies grown from puromycin selected CRISPR knock-in pools were assessed for incorporation of puromycin resistance vector via PCR amplification using primers flanking incorporation site.

Supplemental Figure 1

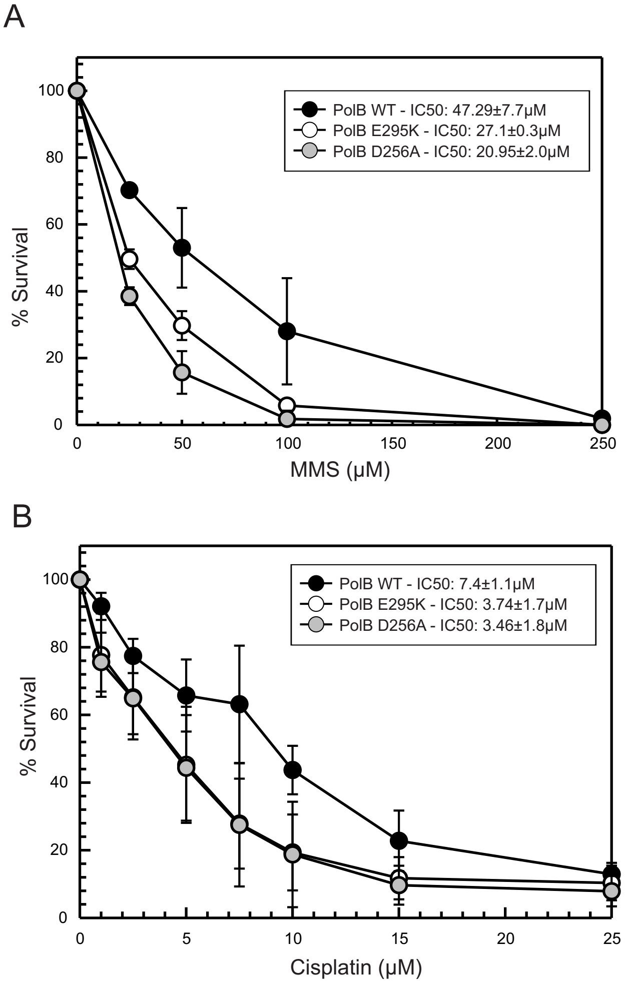

Supplemental Figure 2.

A

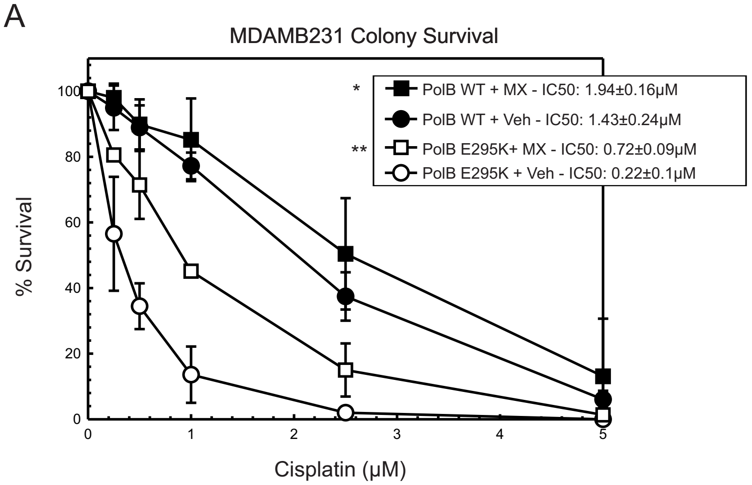


B

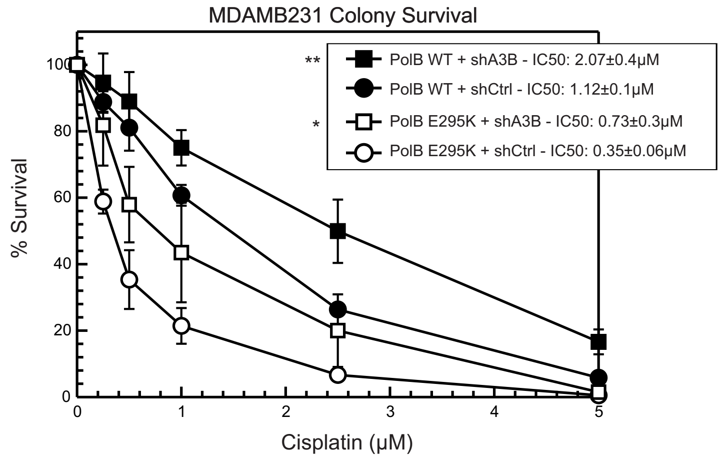


C

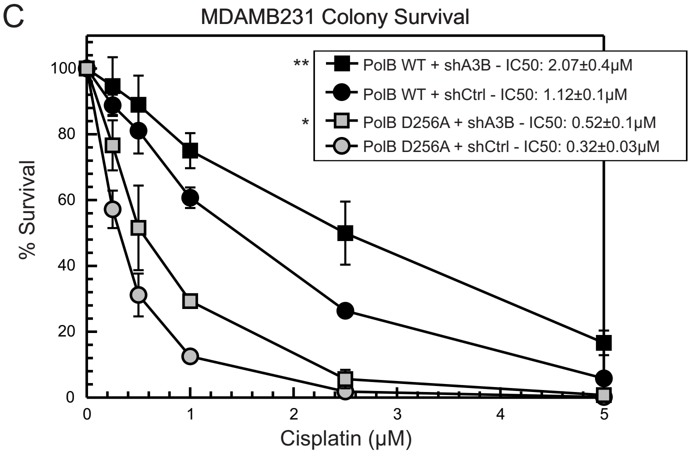

Supplemental Figure 3

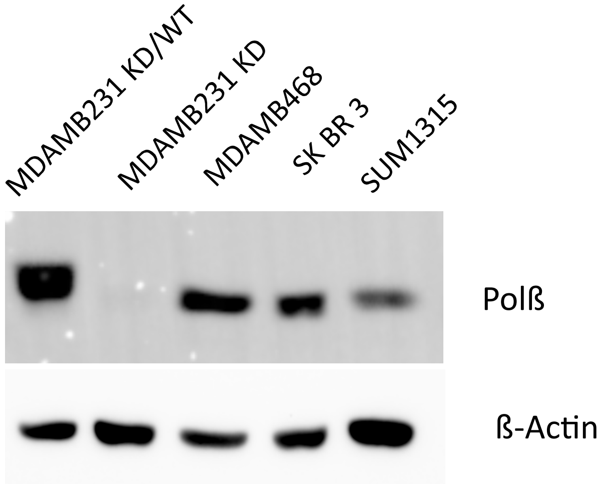

Supplemental Figure 4

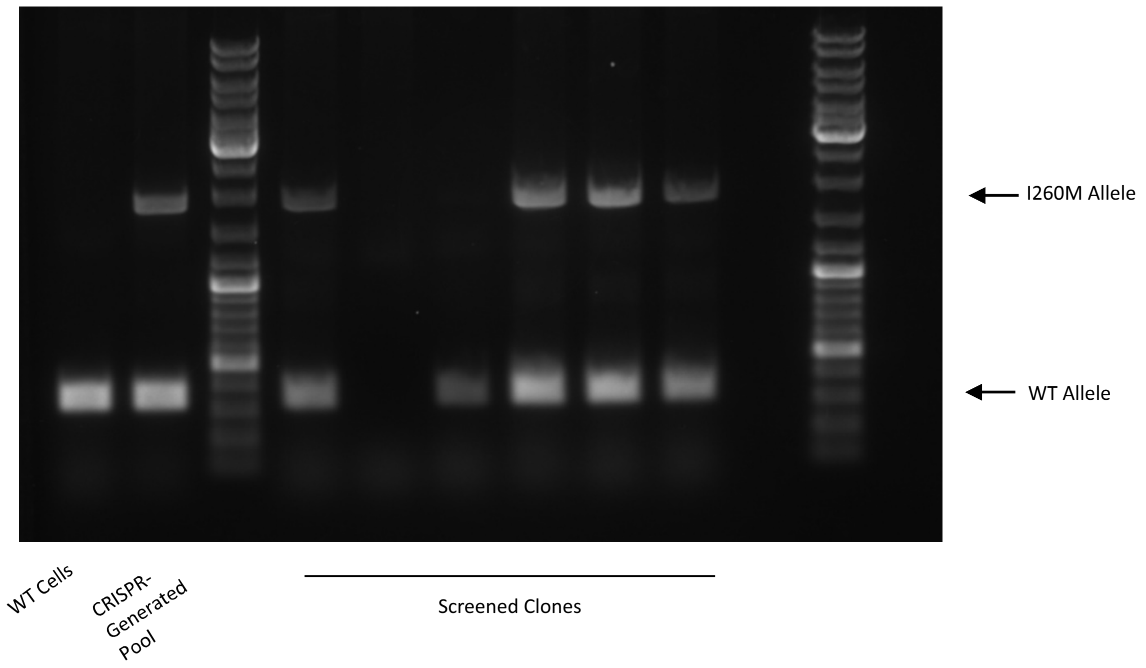
